# Defining the genes required for survival of Mycobacterium bovis in the bovine host offers novel insights into the genetic basis of survival of pathogenic mycobacteria

**DOI:** 10.1101/2022.03.15.484275

**Authors:** Amanda J Gibson, Jennifer Stiens, Ian J Passmore, Valwynne Faulkner, Josephous Miculob, Sam Willcocks, Michael Coad, Stefan Berg, Dirk Werling, Brendan W Wren, Irene Nobeli, Bernardo Villarreal-Ramos, Sharon L Kendall

**Affiliations:** Centre for Emerging, Endemic and Exotic Diseases, Pathobiology and Population Sciences, Royal Veterinary College, Hawkshead Lane, North Mymms, Hatfield, AL9 7TA, United Kingdom.; Institute of Structural and Molecular Biology, Biological Sciences, Birkbeck, University of London, Malet Street, London, WC1E 7HX, United Kingdom.; London School of Hygiene and Tropical Medicine, Keppel Street, London, WC1E 7HT, United Kingdom.; Animal and Plant Health Agency, Woodham Ln, Addlestone, Surrey, KT15 3NB, United Kingdom; Centre of Excellence for Bovine Tuberculosis, IBERS, Aberystwyth University, Penglais, Aberystwyth, Ceredigion, SY23 3EE, United Kingdom; Systems Chemical Biology of Infection and Resistance Laboratory, The Francis Crick Institute, 1 Midland Road, London 15 NW1 1AT, United Kingdom.

## Abstract

Tuberculosis has severe impacts in both humans and animals. Understanding the genetic basis of survival of both *Mycobacterium tuberculosis*, the human adapted species, and *Mycobacterium bovis*, the animal adapted species is crucial to deciphering the biology of both pathogens. There are several studies that identify the genes required for survival of *M. tuberculosis in vivo* using mouse models, however, there are currently no studies probing the genetic basis of survival of *M. bovis in vivo.* In this study we utilise transposon insertion sequencing in *M. bovis* to determine the genes required for survival in cattle. We identify genes encoding established mycobacterial virulence functions such as the ESX-1 secretion system, PDIM synthesis, mycobactin synthesis and cholesterol catabolism that are required *in vivo*. We show that, as in *M. tuberculosis, phoPR* is required by *M. bovis in vivo* despite the known defect in signalling through this system. Comparison to studies performed in glycerol adapted species such as *M. bovis* BCG and *M. tuberculosis* suggests that there are differences in the requirement for genes involved in cholesterol import (*mce4* operon), oxidation (*hsd*) and detoxification (*cyp125*). We report good correlation with existing mycobacterial virulence functions, but also find several novel virulence factors, including genes involved in protein mannosylation, aspartate metabolism and glycerol-phosphate metabolism. These findings further extend our knowledge of the genetic basis of survival *in vivo* in bacteria that cause tuberculosis and provide insight for the development of novel diagnostics and therapeutics.

**Importance:** This is the first report of the genetic requirements of an animal adapted member of the MTBC in a natural host. *M. bovis* has devastating impacts in cattle and bovine tuberculosis is a considerable economic, animal welfare and public health concern. The data highlight the importance of mycobacterial cholesterol catabolism and identifies several new virulence factors. Additionally, the work informs the development of novel differential diagnostics and therapeutics for TB in both human and animal populations.

## Introduction

Bacteria belonging to the *Mycobacterium tuberculosis* complex (MTBC) have devastating impacts in both animal and human populations. *Mycobacterium bovis,* an animal adapted member of the MTBC and one of the main causative agents of bovine tuberculosis (bTB), remains endemic in some high-income settings despite the implementation of a test and slaughter policy. In low- and middle-income settings, the presence of bTB in livestock combined with the absence of rigorous control measures contributes to the risk of zoonotic transmission (1, 2). Control measures based on cattle vaccination utilise the live attenuated vaccine *M. bovis* BCG but the efficacy of this vaccine still remains low in field situations (3, 4). In addition to vaccines, the development of diagnostic tools for the identification of infected individuals is crucial for the management of transmission. Currently, vaccination with *M. bovis* BCG sensitises animals to the diagnostic tuberculin skin test, therefore, sensitive and specific differentiating diagnostic strategies are a current imperative (5, 6).

The increased accessibility of whole genome fitness screens has allowed the assessment of the impacts of the loss of gene function on bacterial survival (7). Such screens have been invaluable in identifying novel drug targets or candidates for the generation of new live attenuated vaccines in a number of bacterial pathogens, including *M. tuberculosis* (8–13). Studies utilising whole genome transposon mutagenesis screens to examine gene fitness *in vivo* in *M. tuberculosis* have been limited to mouse models (8, 9, 13). These models do not faithfully replicate the granulomatous pathology associated with TB, nor do mice contain the same repertoire of CD1 molecules expressed by bovine T cells required to present mycobacterial lipid antigens (14). Whole genome transposon mutagenesis screens utilising non-human primates are limited because screening is restricted to smaller mutant pools (15). To date, transposon insertion sequencing (Tn-seq) based studies in the context of bTB in cattle have only been performed using *M. bovis* BCG strains (16, 17).

In this study we use Tn-seq to determine the genes required for survival of *M. bovis* directly in cattle. We show that genes involved in the biosynthesis of phthiocerol dimycocerosates (PDIMs), the ESX-1 secretion system, cholesterol catabolism, and mycobactin biosynthesis are essential for survival in cattle, corroborating current knowledge of gene essentiality in members of the MTBC (8, 9, 13, 16, 17). We identify differences in the requirement for genes involved in cholesterol transport and oxidation in the fully virulent *M. bovis* strain. We also identify several novel genes required for survival *in vivo* that have not been previously described in members of the MTBC.

## Results and Discussion

### Generation and sequencing of the input library

We generated a transposon library in *M. bovis* AF2122/97 using the MycomarT7 phagemid system as previously described (18, 19). Sequencing of the input library showed that transposon insertions were evenly distributed around the genome and 27,751 of the possible 73,536 thymine–adenine dinucleotide (TA) sites contained an insertion representing a saturation density of ∼38% (Supplementary Figure S1 and Supplementary Table S1-input library). The *M. bovis* AF2122/97 genome has 3,989 coding sequences and insertions were obtained in 3,319 of these, therefore the input library contained insertions in 83% of the total coding sequences.

### Mycobacterium bovis specific immune responses were observed in cattle

Twenty-four clinically healthy calves of approximately 6 months of age were inoculated with the library through the endobronchial route. Infection was monitored by IFN-γ release assay (IGRA) at the time of inoculation and 2 weeks post infection. *M. bovis* specific immune responses were observed for all study animals at 2 weeks post infection (Figure 1A and B). Each animal presented very low background of circulating IFN-γ together with a statistically significant increase in IFN-γ release in response to PPD-B compared to PPD-A antigens (Figure 1C; *** p ≤ 0.001). This indicates that infection with the library was successfully established in the cattle.

**Figure 1.**
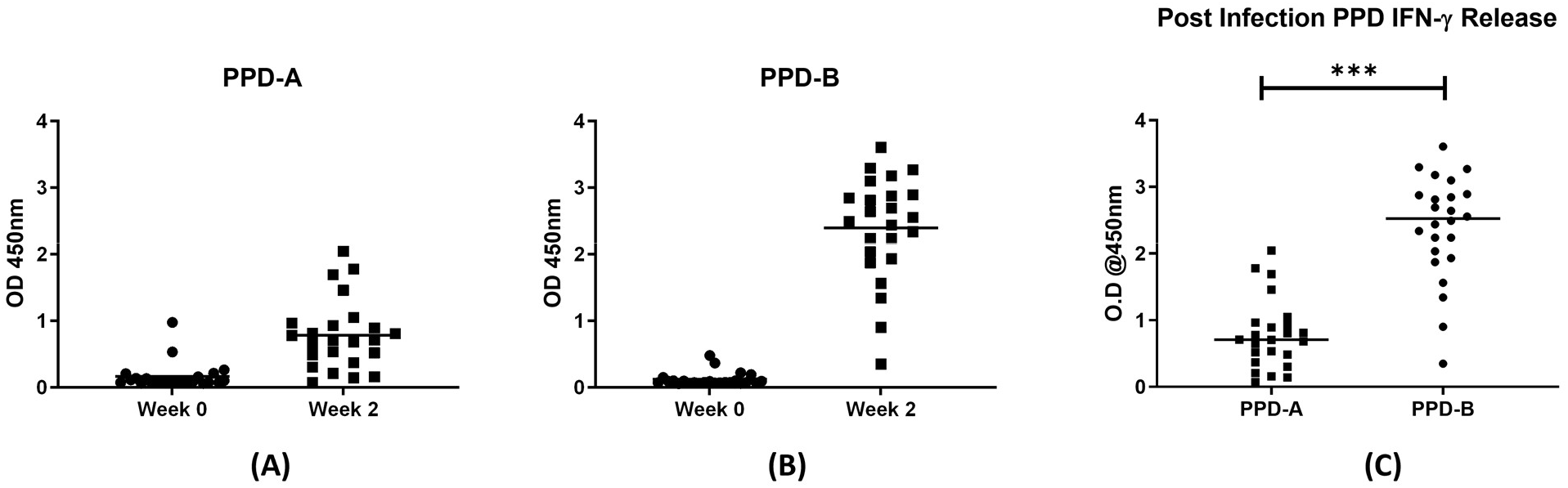
bTB specific IFN-gamma release in cattle infected with the *M. bovis* Tn-library. Blood was collected from all 24 animals on the day of infection and 2 weeks later. No response was detected to either PPD-A or PPD-B antigen stimulation prior to infection (Figure 1A and Figure 1B, week 0). All animals presented a significant and specific response to PPD-B compared to PPD-A as determined by a paired T-test using GraphPad Prism (Figure 1C). *** p ≤ 0.001

### Pathology associated with infection was greater in the lung and thoracic lymph nodes

Animals were culled at 6 weeks post infection. Lung sections and upper (head and neck) and lower (thoracic) respiratory tract associated lymph nodes were examined for gross lesions. Lesions typical of *M. bovis* infection were observed in the tissues examined. Pathology scores are shown in Figure 2A. Greater pathology was observed in lung and thoracic lymph nodes compared to the head and neck lymph nodes.

**Figure 2.**
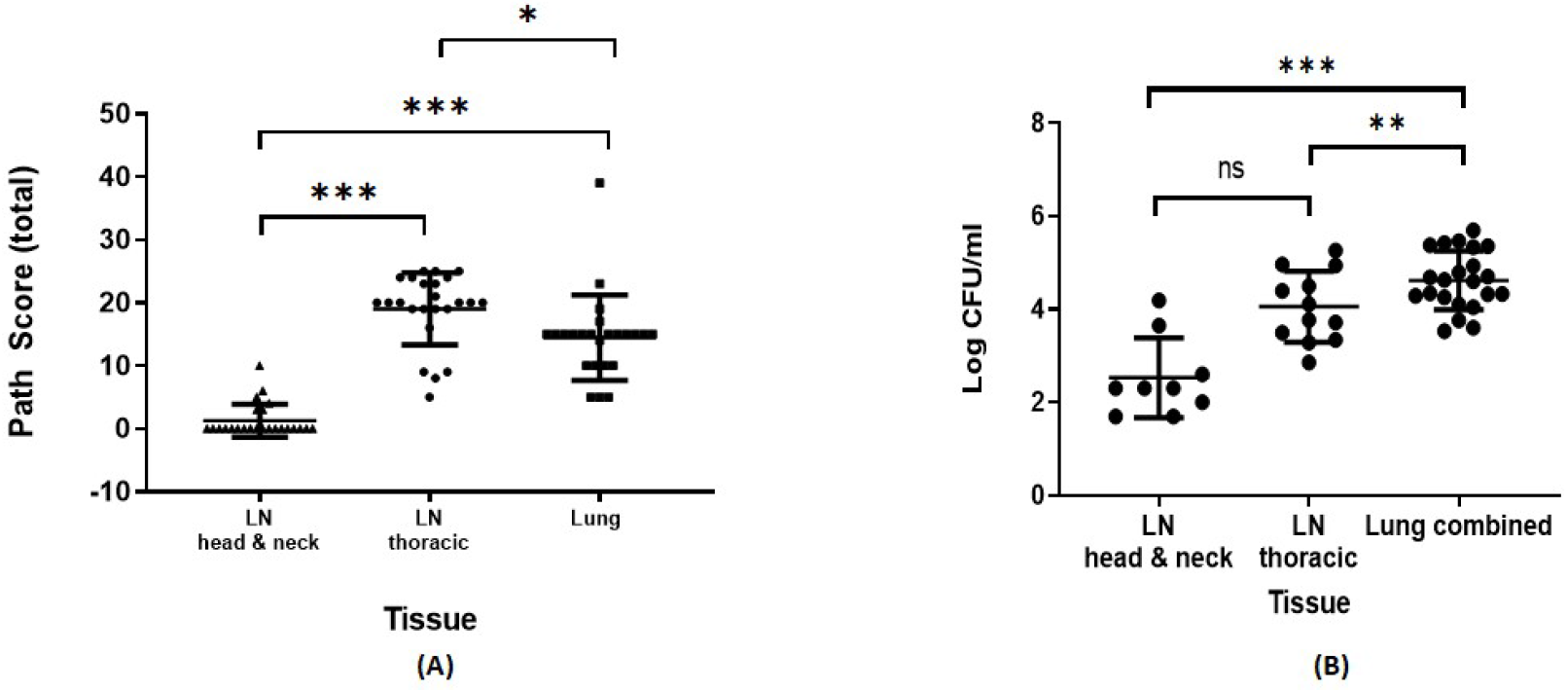
Tissue pathology and bacterial load in tissue sites. Six weeks after infection animals were subjected to post-mortem examination. Gross pathology and evidence of TB-like granulomas lesions were scored. Data presented is the mean across animals of the total scores for each tissue group from 24 animals +/- the standard deviation. Lung and thoracic lymph nodes were observed to contain the highest pathology compared to head and neck lymph nodes (Figure 2A). For bacterial load estimation, aliquots of macerates were plated onto modified 7H11 agar containing kanamycin. Colonies were counted after 3-4 weeks growth. Data are presented as mean CFU/ml per collected tissue group +/- standard deviation. Lung tissue contained the highest bacterial burden compared to thoracic and head and neck lymph nodes as determined by one-way ANOVA analysis using GraphPad Prism (Figure 2b). *** p ≤ 0.001, ** p = 0.002, *p=0.01

### Higher bacterial loads were associated with the lung and thoracic lymph nodes

Bacterial counts were highest in lesions derived from the lung compared to those from the thoracic lymph nodes and head and neck lymph nodes (Figure 2B). The lowest bacterial counts were observed within the head and neck lymph nodes. However, this was not significant when compared to thoracic lymph nodes. The volume of each macerate varied depending on lesion size. Considering macerate volume, average bacterial loads of 10^7^, 10^6^ and 10^5^ were recovered from lesions from samples of the lungs, thoracic lymph nodes and head and neck lymph nodes, respectively.

### Recovery and sequencing of in vivo selected transposon libraries

In order to recover the Tn library from harvested tissue ∼ 10^5 -^10^6^ CFU from lungs and thoracic lymph nodes were plated onto several 140 mm modified 7H11 plates containing kanamycin to minimize competition between mutants. Samples from 4 cattle were lost due to fungal contamination, therefore the samples processed represent samples from 20 cattle. Lung samples were plated from all 20 animals and thoracic lymph nodes samples were plated from 6 cattle. Bacteria were grown for 4-6 weeks before harvesting for genomic DNA extraction and subsequent sequencing (see Supplementary Table S1 for assignation of sequencing files to samples). The diversities of the output libraries were compared to the input library for each sample (Supplementary Figure S2 and Table S1). On average, libraries recovered from lung lesions from 20 different cattle contained 14,456 unique mutants and those recovered from the thoracic lymph nodes contained an average of 16,210 unique mutants. Given that the input library contained 27,751 unique mutants this represented a loss of diversity of ∼ 40- 50%. Good coverage of coding sequences (CDSs) was maintained as the output libraries still contained insertions in (on average) 68-70% of the open reading frames.

Comparison of the read counts between the input and output libraries allowed a measurement of the impact of the insertion on the survival of mutants in cattle. The results are represented as a mean log_2_ fold-change in the output compared with the input for each gene. The entire dataset is shown in supplementary Table S4 and a volcano plot from the lungs and thoracic lymph node of two representative animals is shown in Supplementary Figure S3. Comparison of the mean log_2_ fold-change between lung and lymph node samples showed good correlation (Spearman’s rho = 0.88, p-value <2.2e-16) (Supplementary Figure S4). TRANSIT resampling was performed to compare the composition of the mutant population in the lungs and thoracic lymph nodes of paired cattle, it was also applied to compare all the thoracic lymph nodes with the lungs of all cattle samples. No statistically significant differences were observed indicating that there were no differences in mutant composition between the tissue sites.

No insertion mutants were significantly over-represented in the output library in any of the animals. Although interestingly, insertions in *MB0025,* a gene that is unique to *M. bovis* appeared to improve growth in cattle as mean log_2_ fold-changes of +3.9 (lungs) and +4.2 (lymph nodes) were observed; however, significance criteria were not met in any of the animals. In order to define a list of attenuating mutations, we used a similar approach to that used in a previous study with an *M. bovis* BCG library in cattle (16). Insertions in genes were defined as attenuating if they had log_2_ fold-change of -1.5 or below and an adjusted p-value of <0.05 in at least half of the animals (Table S4, significant in 50% of cattle tab). When using these criteria, there were 141 genes where insertions caused significant attenuation in the lungs or the thoracic lymph nodes, 20 genes that reached significance only in the lungs (shown in red) and 16 genes that reached significance only in the thoracic lymph nodes (shown in green). Of the 141 genes, 109 had been previously described as being required *in vivo* in *M. tuberculosis* H37Rv in mouse models through the use of whole genome Tn screens representing ∼77% overlap with the previous literature (8, 9, 13).

### Comparison with mutations known to cause attenuation in the MTBC

Insertions in the RD1 encoded ESX-1 type VII secretion system secreting virulence factors and immunodominant antigens EsxA (CFP-10) and EsxB (ESAT-6) are expected to cause attenuation (20). The impacts of insertions in this region are summarised in Figure 3 but are also available in Supplementary Table S4 (RD regions tab) and Supplementary Figure S5. Insertions in genes encoding the structural components of the apparatus (*eccB1, eccCa1, eccCb1, eccD1*) were severely attenuating (log_2_ fold-change -6 to -9). Insertions in *eccA1,* which also codes for a structural component of the apparatus, were less impactful (log_2_ fold-changes of -2 to -3) despite good insertion saturation in this gene. This is supported by the work of others who have shown that deletion of *eccA1* in *Mycobacterium marinum* leads to only a partial secretion defect (21). There were no impacts seen due to insertions in accessory genes *espJ, espK* and *espH.* The lack of attenuation seen in *espK* mutants is supported by other studies showing that this gene is dispensable for secretion through the apparatus and is not required for virulence of *M. bovis* in guinea pigs (22, 23). Insertions in *esxA* and *esxB* resulted in severe attenuation (log_2_ fold-change of -6) but this did not reach significance cut-offs (adj. p=<0.05) in any of the cattle. This is likely to be due to the small number of TAs in these genes which makes it challenging to measure mutant frequency.

**Figure 3.**
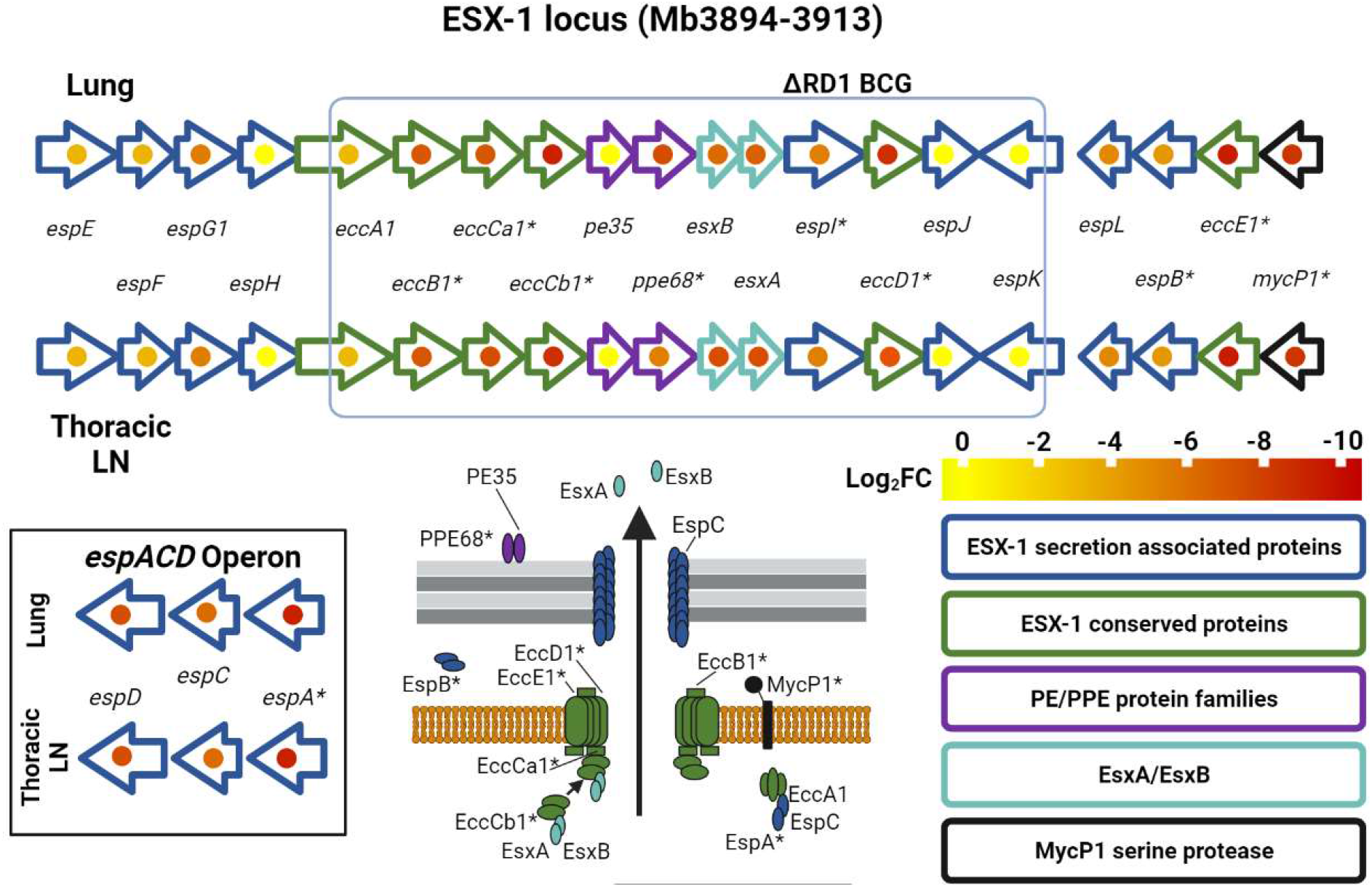
Fold-changes caused by transposon insertions in the ESX-1 secretion system in the lungs and lymph nodes of infected cattle. Asterisks indicate that genes had an adjusted p-value of <0.05 in at least half of the animals. The genes are grouped according to function as indicated by the colour scheme. The log_2_ fold-change are indicated on a yellow to red scale and present as a dot in the centre of the gene.

The highest levels of attenuation seen were in genes involved in the synthesis of the cell wall virulence lipids PDIMs (*ppsABCDE* and *mas* with log_2_ fold-changes of ∼-10 commonly seen). PDIM synthesis is well known to be required for the survival of *M. tuberculosis* and *M. bovis* in mice and guinea pigs (24, 25). Insertions in genes involved in the synthesis of PDIMs were the most under-represented (log_2_ fold-changes of -8 to -10) in the output library (Figure 4, Supplementary Table S4, mycolipids tab). MmpL7 is involved in PDIM transport and there is evidence that it is phosphorylated by the serine-threonine kinase PknD (26). PknD-MmpL7 interactions are thought to be perturbed in *M. bovis* as *pknD* is split into two coding sequences in the bovine pathogen by a frameshift mutation (27). The data presented here suggest that MmpL7 still functions despite the frameshift mutation.

**Figure 4.**
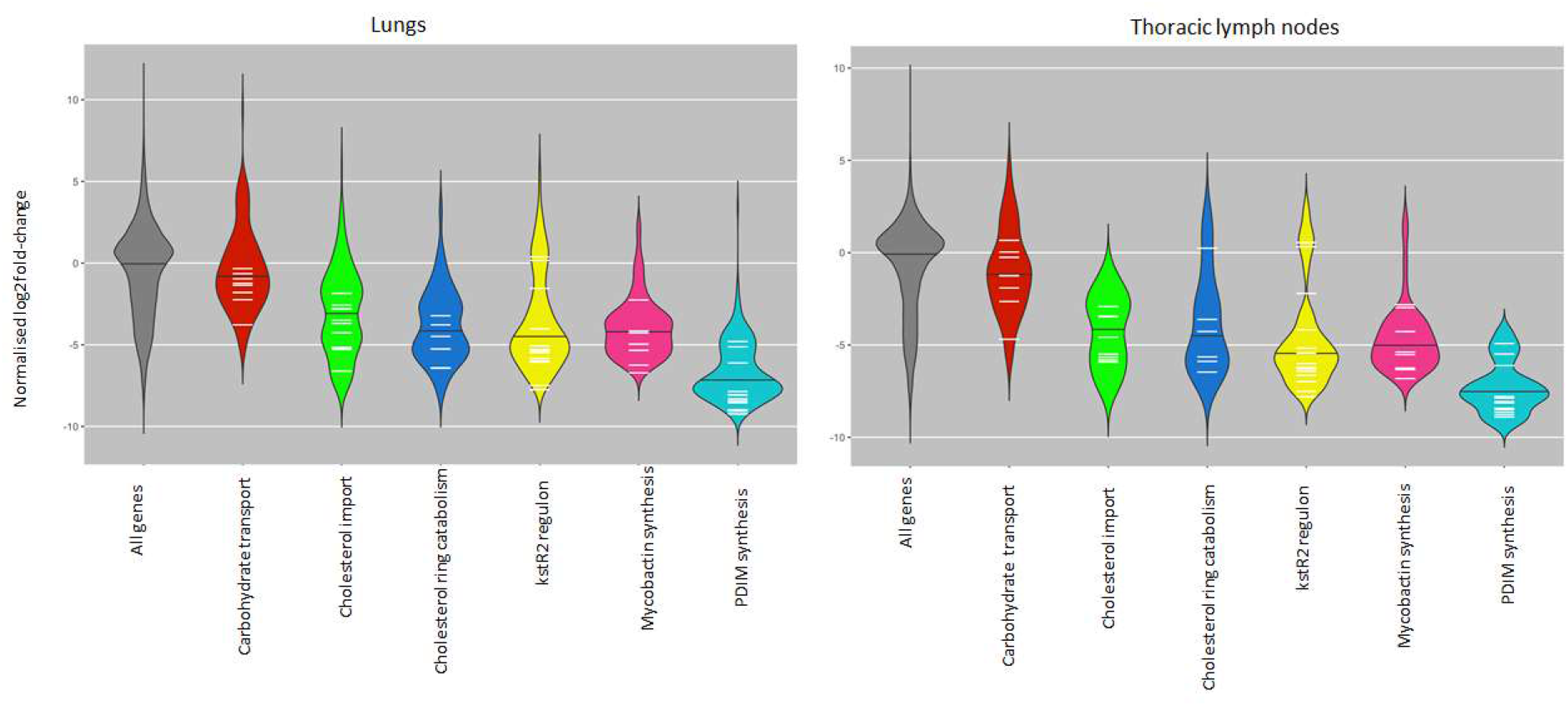
Violin plot of normalised log_2_ fold changes in gene insertions recovered from bovine lung or thoracic lymph node tissue samples in selected gene groups. Black bars indicate overall median of normalized log_2_ fold-change among genes in grouping. White bars indicate mean log_2_ fold-change for each gene in the group across all samples in either lung or lymph node tissue

Iron restriction is thought to be a mechanism by which the host responds to mycobacterial infection, although different cellular compartments may be more restrictive than others (28). Insertion in many of the genes involved in mycobactin synthesis *(Mb2406-Mb2398, mbtJ-mbtH)* were attenuating in cattle (Figure 4, Supplementary Table S4, mycobactin synthesis tab). As mycobactin is required for the acquisition of iron, this confirms that, like other members of the MTBC, needs to scavenge iron from the host for survival (13, 16).

The role of the cholesterol catabolism in *M. tuberculosis* is well documented and it is required for both energy generation and manipulation of the immune response (29–31). Cholesterol uptake is mediated by the Mce4 transporter coded by the *mce4* operon *Rv3492c-Rv3501c* (*Mb3522c-MB3531c*) (32, 33). It has been suggested that an alternative cholesterol acquisition pathway operates in *M. bovis* BCG Danish as, unlike insertions in genes in the down-stream catabolic pathway, insertions in the *mce4* operon do not result in attenuation in this strain (16). In contrast, our study shows that cholesterol transport via the Mce4 transporter is required in *M. bovis* (Figure 4, Supplementary Table S4 -cholesterol catabolism tab, Figure 5). This corroborates work performed in *M. tuberculosis*, where Mce4 has been shown to be required for growth in chronically infected mice (9, 32). Propionyl-coA generated from the catabolism of cholesterol is toxic and detoxification mechanisms include incorporation into PDIMs (34, 35). The observation that BCG Danish contains a lower amount of PDIMs compared to BCG Pasteur (16) suggests a correlation between Mce4 mediated cholesterol transport and PDIM synthesis and previous studies have demonstrated an increase in PDIM biosynthesis as a result of *mce4* over-expression (36). PDIMs biosynthesis genes are over-expressed in *M. bovis* compared to *M. tuberculosis* (27) and comparison of our dataset with Tn-seq studies performed in *M. tuberculosis* (9) indicates an over-reliance of *M. bovis* on cholesterol transport through the Mce4 transporter (Figure 5).

**Figure 5.**
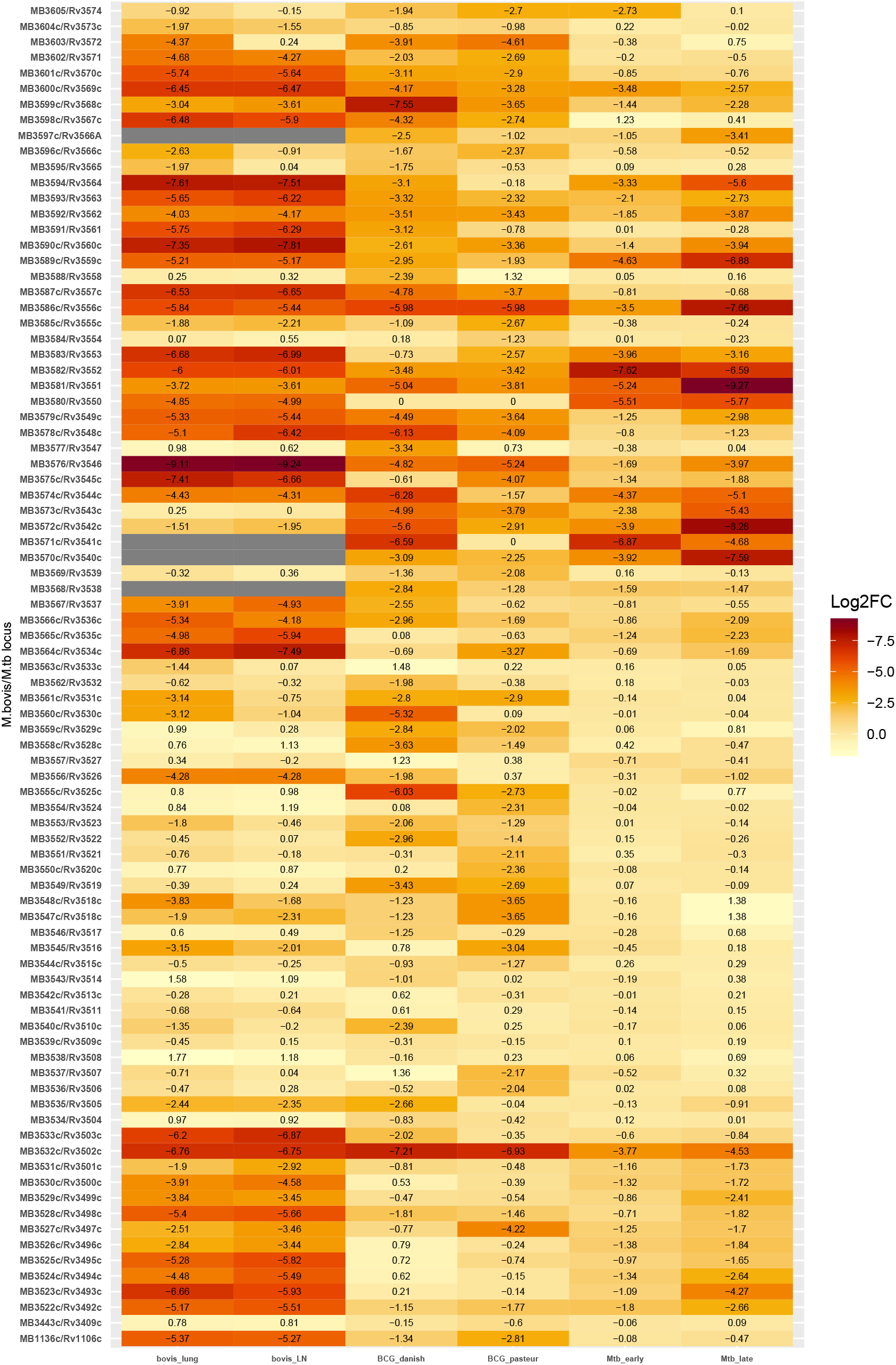
Comparison of reported log_2_ fold-change in *M. bovis*, *M. bovis* BCG and Mtb transposon insertion sequencing experiments for orthologous genes in the cholesterol catabolic pathway. Greatest attenuation (most negative log_2_ fold-change) is coloured by darkest red. Studies used for comparison include Mendum et al., (24) and Bellarose et al., (9). Grey bars represent genes for which there is no information as they were either ES or GD in input library or had less than 5 insertions in any TA site in any sample (input and all output).

Early stages of cholesterol catabolism involve the oxidation of cholesterol to cholestenone, a reaction catalysed by the 3β-hydroxysteroid dehydrogenase (*hsd*) encoded by *Rv1106c/Mb1136c* (37). The cytochrome P450 Cyp125 (*Mb3575c/Rv3545c*) is required for the subsequent detoxification of cholestenone (38). Insertions in both *hsd* and *cyp125* in *M. bovis* were severely attenuating with log_2_ fold-changes of ∼-5 to -7 (Supplementary Table S4 - cholesterol catabolism tab, Figure 5). Previous studies have shown that these genes are not required for the survival of *M. tuberculosis* in macrophages or in guinea pigs and this is thought to be due to the availability of other carbon sources, including glycolytic substrates, *in vivo* (37, 39–43). *M. bovis* is more restricted in metabolic capabilities and is unable to generate energy from glycolytic intermediates, largely due to a disrupted pyruvate kinase encoded by *pykA* (44, 45). The essentiality of *hsd* and *cyp125* during infection for *M. bovis* but not *M. tuberculosis* supports the hypothesis of an over-reliance of *M. bovis* on cholesterol. Given the potential for the use of host cholesterol metabolites as diagnostic biomarkers, this observation might have applications in the development of differential diagnostics (46).

### Genes that are differentially expressed between Mycobacterium bovis and Mycobacterium tuberculosis

Several studies have identified key expression differences between *M. bovis* and *M. tuberculosis* (27, 47, 48). We examined the dataset for insights on the role of differentially expressed genes and transcriptional regulators during infection. One important regulatory system in *M. tuberculosis* is the two-component regulatory system PhoPR and deletions in the *phoPR* genes alongside *fadD26* are attenuating mutations in the live vaccine MTBVAC (49–51). Our data show that insertions in both *phoPR* and *fadD26* were severely attenuating with log_2_ fold-changes of -6 to -9 (Figure 6, Supplementary Table S4, *phoPR* regulon tab and mycolipids tab). This reinforces the role of this system in virulence, despite the presence of a single nucleotide polymorphism (SNP) in the sensor kinase *phoR* that impacts signalling through the system in *M. bovis* (52). Signal potentiation via *phoR* is required for secretion of ESAT-6 through the ESX-1 secretory system and *M. bovis* is known to have compensatory mutations elsewhere in the genome, e.g. in the *espACD* operon, that restores ESAT-6 secretion in the face of a deficient signalling system (49, 52, 53). Our data also show that Tn insertions in *espA* of the *espACD* operon (required for ESAT-6 secretion) and in *mprA*, a transcriptional regulator of that operon (54) were severely attenuating (log2 fold-changes -7 to -9), emphasising the relevance of ESAT-6 as a virulence factor.

**Figure 6.**
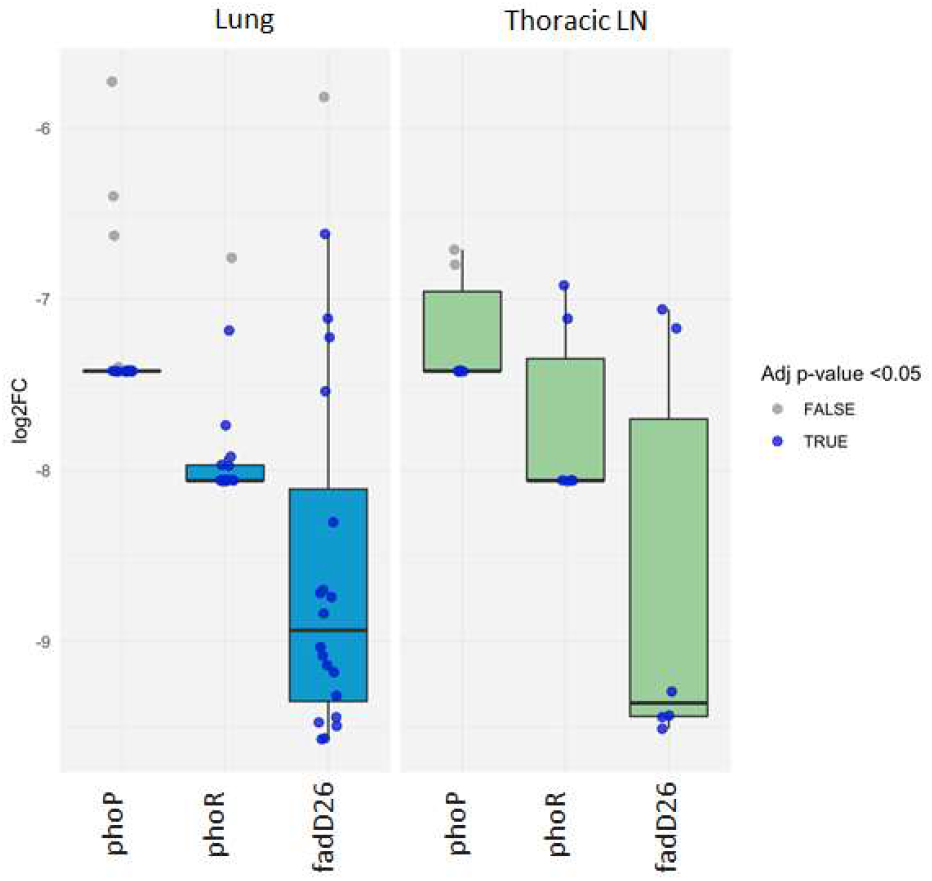
Fold-changes caused by transposon insertions in phoP, phoR and fadD26 in the lungs and lymph nodes of infected cattle. Samples with adjusted p-values (BH-fdr corrected) < 0.05 are indicated with purple points.

Studies comparing differences in expression during *in vitro* growth between *M. bovis* and *M. tuberculosis* show that genes involved in sulfolipid (SL-1) biosynthesis are expressed at lower levels in *M. bovis* compared to *M. tuberculosis* (27, 47). Interestingly, insertions in genes involved in SL-1 biosynthesis (*Mb3850-Mb3856*) are not attenuating *in vivo* (Supplementary File S4, mycolipids tab), reinforcing the lack of importance of SL-1 for *M. bovis in vivo,* at least at the stages of infection studied here.

One of the most highly attenuating insertions occurred in *Mb0222/Rv0216* (log_2_ fold change –8 to -9). This gene has been shown to be highly (> 10-fold) over-expressed in *M. bovis* compared with *M. tuberculosis* but the physiological function of this gene is not currently known. The secreted antigens MPB70 and MPB83, encoded by *Mb2900* and *Mb2898* are also over-expressed in *M. bovis* and play a role in host-specific immune responses, however, insertions in these genes did not cause attenuation *in vivo* in our dataset (55).

### Novel attenuating mutations

We identified 32 genes that were required for survival of *M. bovis* in cattle that had not been previously described as being essential *in vivo* through transposon mutagenesis screens of *M. tuberculosis* in mouse models (8, 9, 13) (see Supplementary Table 4, Significant in 50% of cattle tab). While writing this publication, a large scale Tn-seq study that utilised over 120 *M. tuberculosis* libraries and several diverse mouse genotypes (the collaborative cross mouse panel (56)) showed that the panel of genes required for the survival of *M. tuberculosis in vivo* is much larger than previously reported (57). A direct comparison of our dataset with the study by Smith et al., revealed that a further 13 genes were shown to be required in at least one mouse strain in that study. A summary set of the remaining 19 genes is given in Supplementary Table 4, Not in Mtb Tn-seqs tab. Some of these genes have been shown to be attenuated in the mouse model in *M. tuberculosis* through the use of single mutants (58–61).

Included in this list are genes required for phenolic glycolipid synthesis (Figure 7). Insertions in *Mb2971c/Rv2947c* (*pks15/1*) and in *Mb2972c/Rv2948c* (*fadD22*) were attenuating in *M. bovis* but these genes are not required *in vivo* in *M. tuberculosis,* including in the extended panel of mouse genotypes (8, 9, 13, 57). Both *pks15/1* and *fadD22* are involved in the early stages of synthesis of phenolic glycolipids (PGLs) and are involved in virulence (62). The requirement for these genes in *M. bovis* but not in *M. tuberculosis* is consistent with the observation that the Tn-seq studies in *M. tuberculosis* are often carried out using lineage 4 strains (H37Rv and CDC1551) that harbour a frameshift mutation in the *pks15/1* gene, which renders them unable to synthesise PGLs. This removes the requirement for these genes *in vivo* in lineage 4 strains of *M. tuberculosis. pks15/1* has been previously reported to be required for survival of *M. bovis* in a guinea pig model of infection (63).

**Figure 7.**
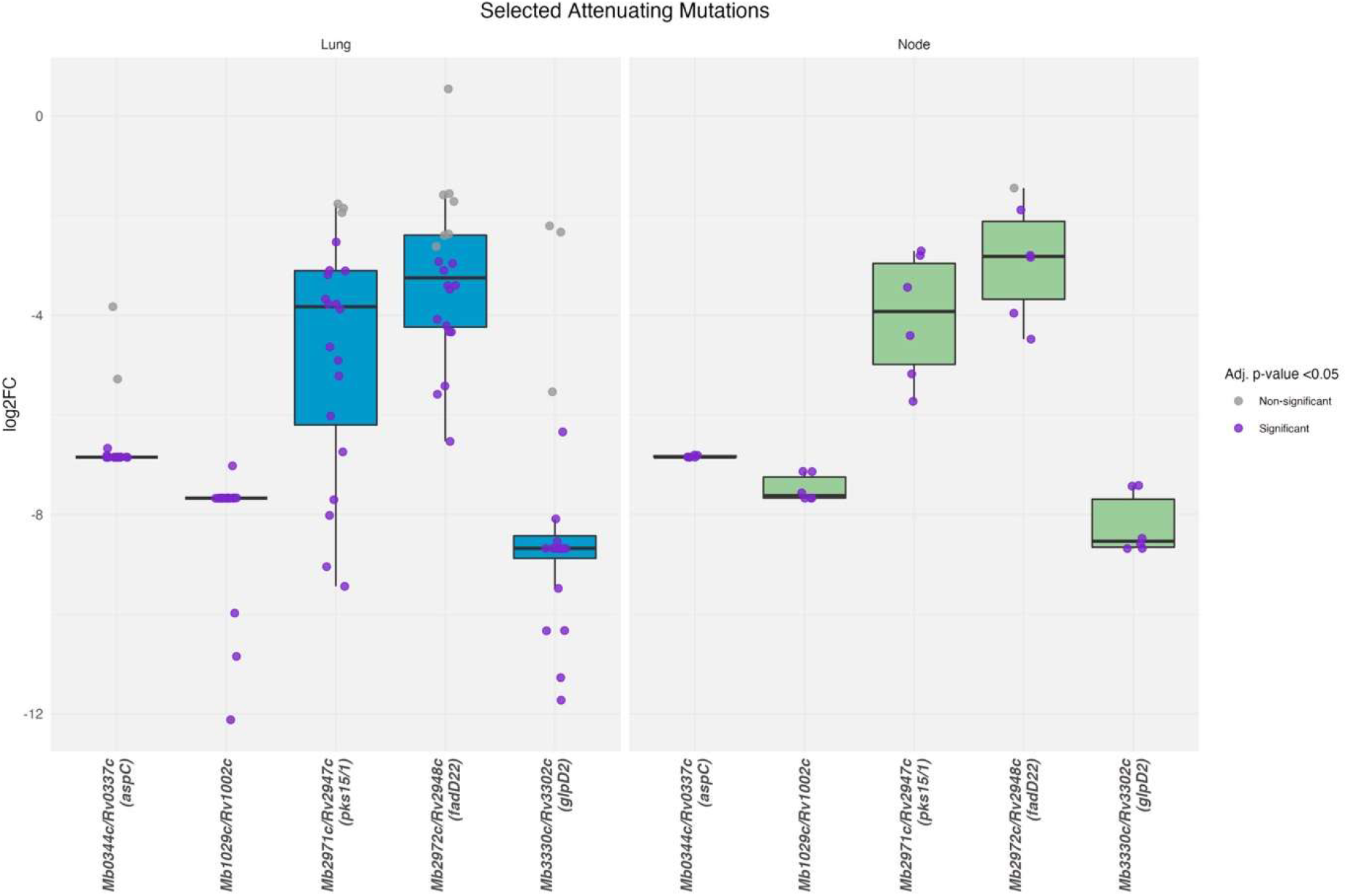
Fold-changes caused by transposon insertions in *pks15/1*, *fadD22*, *Rv1002c*, *aspC* and *glpD2* in the lungs and lymph nodes of infected cattle. Samples with adjusted p-values (BH-fdr corrected) < 0.05 are indicated with purple points.

The list also includes genes that are involved in post-translational modifications such as glycosylation. *Rv1002c* is thought to add mannose groups to secreted proteins and over-expression of this protein in *M. smegmatis* was recently shown to enhance survival *in vivo* and inhibit pro-inflammatory cytokine production (64). The substrates of this protein mannosyltransferase are thought to be several secreted lipoproteins, including LpqW which is involved in the insertion of the virulence lipid LAM at the mycobacterial cell surface (64, 65).

Finally, this list includes two genes (*aspC* and *glpD2*) that are essential *in vitro* in *M. tuberculosis* but not in *M. bovis* (10, 11, 18, 66). Information regarding *aspC* and *glpD2* from Tn-seq approaches is likely to be lacking in *M. tuberculosis* because Tn mutants will not be represented in the input pool. The absence of insertion mutants in these genes in the most recent large-scale *M. tuberculosis* Tn-seq study supports this (57). One of these genes *MB0344c*/*Rv0337c* (*aspC*) is an aspartate aminotransferase involved in the utilisation of amino acids (aspartate) as a nitrogen source (67). The other gene *Mb3303c*/*Rv3302c* (*glpD2*) is a membrane bound glycerol-phosphate dehydrogenase. In *Escherichia coli*, *glpD2* is an essential enzyme, functioning at the central junction of respiration, glycolysis, and phospholipid biosynthesis and catalyses the oxidation of dihydroxyacetone phosphate (DHAP) from glycerol-3-phosphate resulting in the donation of electrons to the electron transport chain (68). Its essentiality *in vitro* in *M. tuberculosis* might be explained by the usage of glycerol during *in vitro* growth in this species. The contribution of the membrane bound *glpD2* in donation of electrons to the electron transport chain, has been suggested but not yet explored in the MTBC (69). Given the interest in the electron transport chain as a chemotherapeutic target in *M. tuberculosis*, the data presented here suggests that inhibition of *glpD2* might be a fruitful approach in the development of new drugs for the treatment of TB in humans (70). The role of this gene in *M. bovis in vivo* is perhaps surprising, given the disruptions in glycerol phosphate uptake and pathways that phosphorylate glycerol in *M. bovis* AF2122/97 (71). However, *M. tuberculosis* is thought to engage in catabolism of membrane derived glycerophospholipids which may be a potential source of glycerol-3-phosphate in members of the complex (72).

## Materials and Methods

### Bacterial strains and culture methods

*M. bovis* strain AF2122/97 was maintained on modified Middlebrook 7H11 (BD Difco^TM^) medium (73). Liquid cultures of *M. bovis* were grown in Middlebrook 7H9 media (BD Difco^TM^) containing 75 mM sodium pyruvate, 0.05% v/v Tween^®^80 and 10% Middlebrook albumin-dextrose-catalase (ADC) (BBL BD Biosciences). Kanamycin at 25 µg/ml was used for selection where appropriate.

### Generation of input transposon mutant library and preparation of the inoculum

Transposon libraries in *M. bovis* were generated as previously described using the MycomarT7 phagemid system as per Majumdar et al with modifications (19). Approximately 66,000 kanamycin resistant transductants were scraped and homogenised in 7H9 medium and stored frozen at -80°C in 1 ml aliquots. CFU counting was performed on the homogenised culture to inform inoculum dosage.

### Cattle Infection

Experiments were carried out according to the UK Animal (Scientific Procedures) Act 1986 under project license PPL70/7737. Ethical permission was obtained from the APHA Animal Welfare Ethical Review Body (AWERB) (UK Home Office PCD number 70/6905). All animal infections were carried out within the APHA large animal biocontainment level 3 facility. Twenty-four Holstein-Friesian crosses of 6 months of age were sourced from an officially TB-free herd. An infectious dose of 7 x 10^4^ CFU was targeted for the “input” library, allowing each mutant to be represented in the library ∼ 2.5-fold. Retrospective counting of the inoculum revealed the actual inoculum for infection contained 4 x 10^4^ CFU. The inoculum was delivered endobronchially in 2 ml of 7H9 medium. In brief, animals were sedated with xylazine (Rompun® 2%, Bayer, France) according to the manufacturer’s instructions (0.2 mL/100 kg, IV route) prior to the insertion of an endoscope through the nasal cavity into the trachea for delivery of the inoculum through a 1.8 mm internal diameter cannula (Veterinary Endoscopy Services, U.K.) above the bronchial opening to the cardiac lobe and the main bifurcation between left and right lobes.

### Infection Monitoring with the IFN-γ release Assay (IGRA)

Blood was collected by jugular venepuncture from animals on the day of the infectious challenge and two weeks after infection. Heparinized whole blood (250 μl) was incubated with purified protein derivative (PPD) from *M. avium* (PPD-A) or PPD from *M. bovis* (PPD-B) (Prionics™) respectively at 25 IU and 30 IU final. Pokeweed mitogen was used as the positive control at 10 µg/mL and a medium-only negative control. After 24 h incubation in 5% (v/v) CO_2_, 95% humidity, 37 °C atmosphere bloods were centrifuged (400 × *g* for 5 min); 120 µl of supernatant was removed and stored at −80 °C for subsequent IFN-γ quantification using the BOVIGAM® kit (Prionics™) in accordance with the manufacturer’s instructions.

### Collection of tissues and gross pathology scores

Six weeks after the initial infection animals were subjected to post-mortem examination. Initially the experiment was designed with two time points; an early time point (6 weeks) and a later time point of 8 weeks. However, due to the unexpected high-levels of pathology seen at the earlier time-points all animals were culled at 6 weeks. Gross pathology and evidence of TB-like granulomas lesions was scored using a modified methodology to that previously described in (74). Tissue from head and neck lymph nodes (from the right and left sub-mandibular lymph nodes, the right and left medial retropharyngeal lymph nodes), thoracic lymph nodes (the right and left bronchial lymph nodes, the cranial tracheobronchial lymph nodes, the cranial and caudal mediastinal lymph nodes) and from lung lesions, was collected into sterile containers and frozen at −80 °C until further processing. Frozen tissues were thawed and homogenised in PBS using a Seward Stomacher Paddle Blender.

### Recovery of the output transposon mutant library from tissues

Tissue macerates collected from study animals were thawed at room temperature, diluted in PBS and plated on modified 7H11 agar to determine bacterial loads. Colony counts were performed after 3-4 weeks growth. For recovery of the library from tissue macerates ∼10^5^-10^6^ CFU were plated from lung lesions and thoracic lymph node lesions onto modified 7H11 media containing 25 µg/ml kanamycin. The colonies were plated over several 140 mm petri dishes to minimise competition between mutants. The colonies were harvested after 4-6 weeks growth and genomic DNA extracted.

### Genomic DNA extraction

Genomic DNA from the input and recovered libraries was isolated by an extended bead beating procedure with detergent-based lysis, phenol-chloroform DNA extraction and precipitation as previously described (18). DNA quality was assessed by nano-spectrometry (DeNovix) and gel electrophoresis and quantified by Qubit analysis using the Broad Range Assay Kit (ThermoScientific).

### Library preparation for transposon directed insertion sequencing

DNA (2 µg) was resuspended in 50 µL distilled water and sheared to approximately 550 bp fragments using a S220 focussed-ultrasonicator (Covaris), according to the manufacturer’s protocol. Fragmented DNA was repaired using NEBNext blunt-end repair kit (New England Biolabs) and purified using Monarch PCR clean-up kit (NEB). Blunted DNA was A-tailed using NEBNext dA-tailing kit (NEB) and column purified. Custom transposon sequencing adaptors (Supplementary Table S3) were generated by heating an equimolar mix of Com_AdaptorPt1 primer and Com_AdaptorPt2 (P7+index) primers to 95°C for 5 min, followed by cooling by 1°C every 40 s to a final temperature of 4°C in a thermocycler. Adaptors were ligated to A-tailed library fragments using NEBNext quick ligase kit. Transposon-containing fragments were enriched by PCR using ComP7 primer (10 µM) and an equimolar mix of primers P5-IR2a-d primer (10 µM) in a reaction with 50 ng of adaptor ligated template and Phusion DNA polymerase (NEB) in a thermocycler with the following program 98°C 3 min; 4 cycles of 98°C 20s, 70°C 20s, 72°C 1 min; 20 cycles of 98°C 20s, 67°C 20s, 72°C 1 min; 72°C 3 min. Transposon-enriched libraries were subsequently purified with AMPureXP beads (Beckman), pooled together and further purified using AMPure XP beads.

### Data analysis

Indexed libraries were combined, spiked with 20% PhiX, and sequenced on the Illumina Hiseq 3000 platform, using v2 chemistry, generating single-end reads of 250 bp. Raw .fastq sequencing files were analysed for quality and pre-processed using the TRANSIT TPP tool (75) set to default ‘Sassetti’ protocol, in order to remove transposon tags and adapter sequences, and to map reads using BWA-mem to TA sites to the *M. bovis* AF2122/97 genome (NC_002945.3). The TRANSIT ‘tnseq_stats’ tool was run on each sample to assess insertion density, skew, kurtosis and potential amplification bias.

The *M. bovis* AF2122/97 genome was scanned for the non-permissive Himar1 transposon insertion motif (’SGNTANCS’, where S is either G or C and N is any base) as previously described [10]. 6605 sites were identified as non-permissive (approximately 9% of total TA sites) and excluded from resampling analysis. A custom annotation, ‘.prot-table’ for TRANSIT, was created from the *M. bovis* AF2122/97 annotation file (NCBI Accession Number LT708304, version LT708304.1). TRANSIT HMM was run on the input library using the default normalisation (TTR) with LOESS correction for genomic position bias. Each TA site was assigned an essentiality state and genes were assigned an essentiality call based on the assigned state of the TA sites within annotated gene boundaries.

Resampling between the input library and each of the output sample libraries was performed independently using the TRANSIT resampling algorithm and the complete prot-table. TTR normalisation was used for 23 of the samples, and betageom normalisation for the three samples with skew of greater than 50. The initial resampling output files were evaluated to identify genes with very few, or no, reads at any TA site within the gene boundaries in both the input library and output sample libraries. Genes with no read counts greater than 4 at any TA site, in any sample, and with a sum of all reads at any TA site across the 26 samples less than 55, were flagged. Essential and unchanged genes were removed from the prot-table prior to further evaluation. Resampling was further limited to protein-coding genes. Resampling was re-run for each sample using the attenuated prot-table and an edited TRANSIT resampling script to return the left-tail p-value, as the data were expected to reflect attenuation. All p-values were corrected for multiple testing with FDR adjustment.

All analysis and plots were performed using R and R packages, tidyverse and circlize (76–78). Orthologous TB genes were obtained from supplementary data files published by Malone et al, 2018 (27). All scripts, prot-tables and insertion files are available at https://github.com/jenjane118/Mbovis_in-vivo_Tnseq, DOI:10.5281/zenodo.6354151. Sequencing files (.fastq) are deposited in SRA (Bioproject ID: PRJNA816175, Submission ID: SUB11067380)

## Funding

This work was funded by the BBSRC Grant Ref: BB/N004590/1 [awarded to SK (PI), DW (Co-I), BW (Co-I), and SE3314 to BV-R as part of the joint BBSRC-DEFRA EradbTB consortium. AG, IP, and SW were supported by the funding. VF was in receipt of an RVC PhD studentship. AG currently holds a Sêr Cymru II Lectureship funded by the European Research Development Fund and Welsh Government. BV-R is a Ser Cymru II Professor of Immunology at Aberystwyth University. JS is supported by a Bloomsbury Colleges PhD Studentship (LIDo program).

## AUTHOR CONTRIBUTIONS

SK, DW, BW, BV-R and SB undertook funding acquisition and designed the study. AJG, VF, JM, SW, IP, MC carried out the experimental work. Data analysis was done by IN and JS. AJG, JS and SLK wrote the first draft of the manuscript. All authors contributed to the manuscript revision, read, and approved the submitted version.

## Acknowledgements

The authors would like to acknowledge the help and support of APHA colleagues from the BAC4 workgroup and the Pathology Unit, and in particular acknowledge the care and support provided to animals under experimentation by members of APHA’s Animals Sciences Unit. We would like to thank Dr Dany Beste for useful discussions surrounding the role of *glpD2* in mycobacterial metabolism.

## Notes

### Competing Interest Statement

The authors have declared no competing interest.

### Summary of Updates

Improve figure resolution Add another box plot (figure 7)

