## Supplemental Figure S1-S5 for "Defining the genes required for survival of Mycobacterium bovis in the bovine host offers novel insights into the genetic basis of survival of pathogenic mycobacteria"

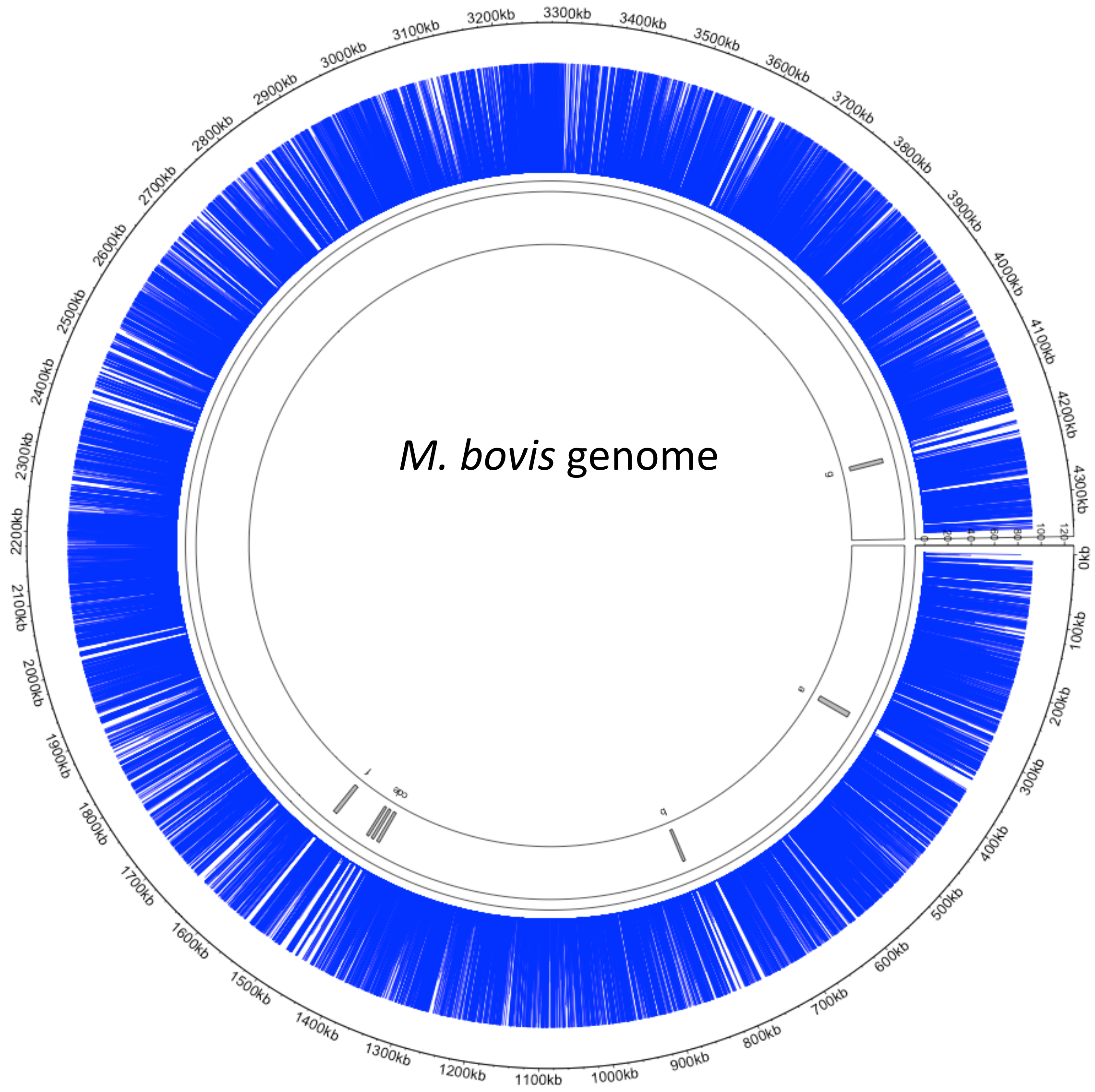


TRANSIT HMM calls for the input library:

| | |
|---|---|
| 528 | Essential |
| 295 | Growth Disadvantage |
| 88 | Growth advantage |
| 3124 | Non-essential |

**Figure S1. Illustration of the transposon insertions around the *M. bovis* genome.** Of the 73,536 possible TA sites 27,751 contain an insertion representing an insertion density of 38% in the input inoculum. The outer ring are the genomic coordinates, the blue lines represent transposon insertions and the gray boxes indicate regions of essential genes that did not have any insertions. Plot made with Circlize (Gu et al, 2014).

Figure S2

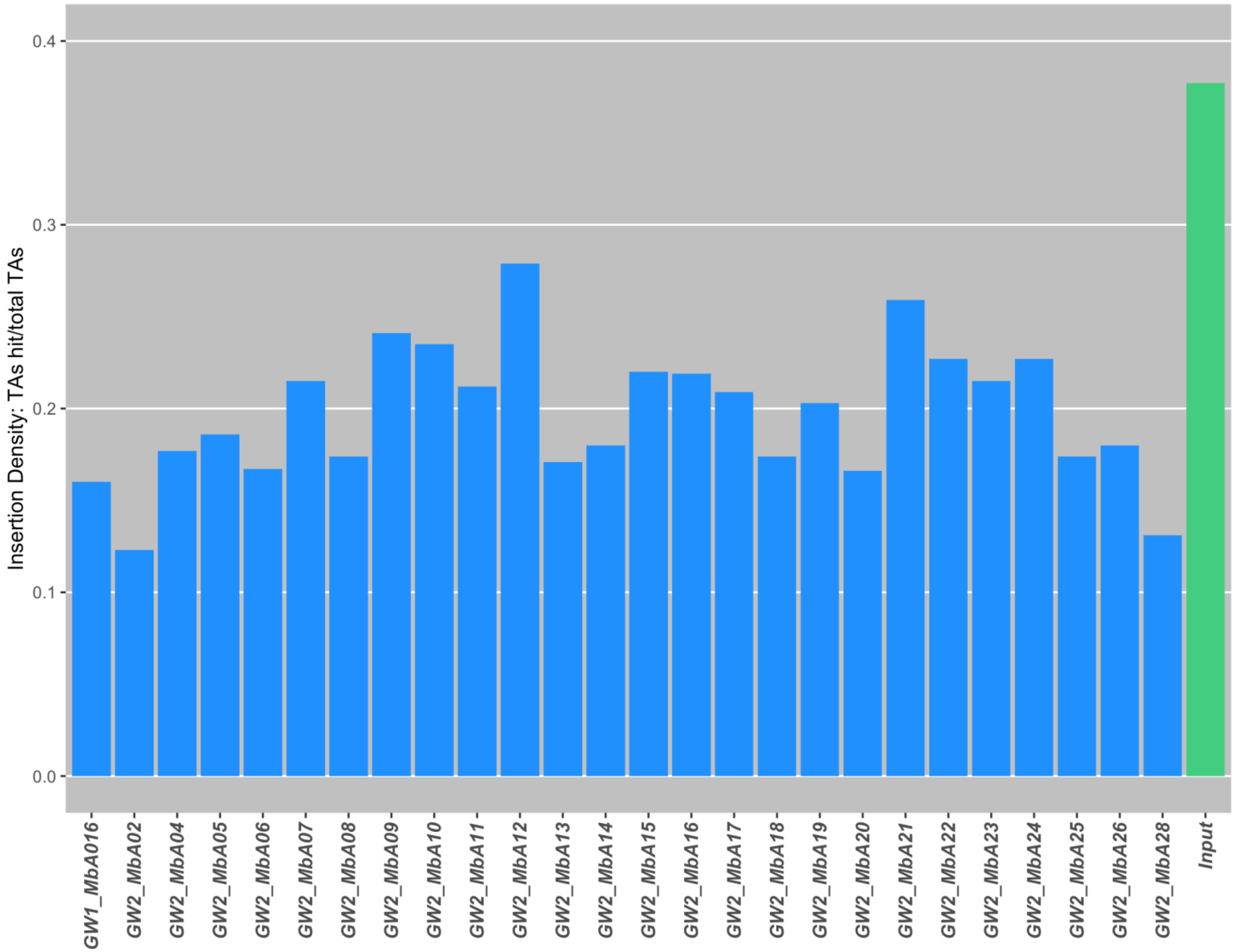


**Figure S2. Diversity of the output library isolated from lung and thoracic lymph node lesions compared to the input library.** On average, libraries recovered from lung lesions contained 14,456 unique mutants and those recovered from the lymph nodes contained an average of 16,210 unique mutants. Insertion density is represented as a proportion of the TA sites that contained insertions. The numbers on the x-axis refer to the sequencing file from that sample and come from individual animals (Bioproject ID: PRJNA816175, Submission ID: SUB11067380).

**Figure S3**

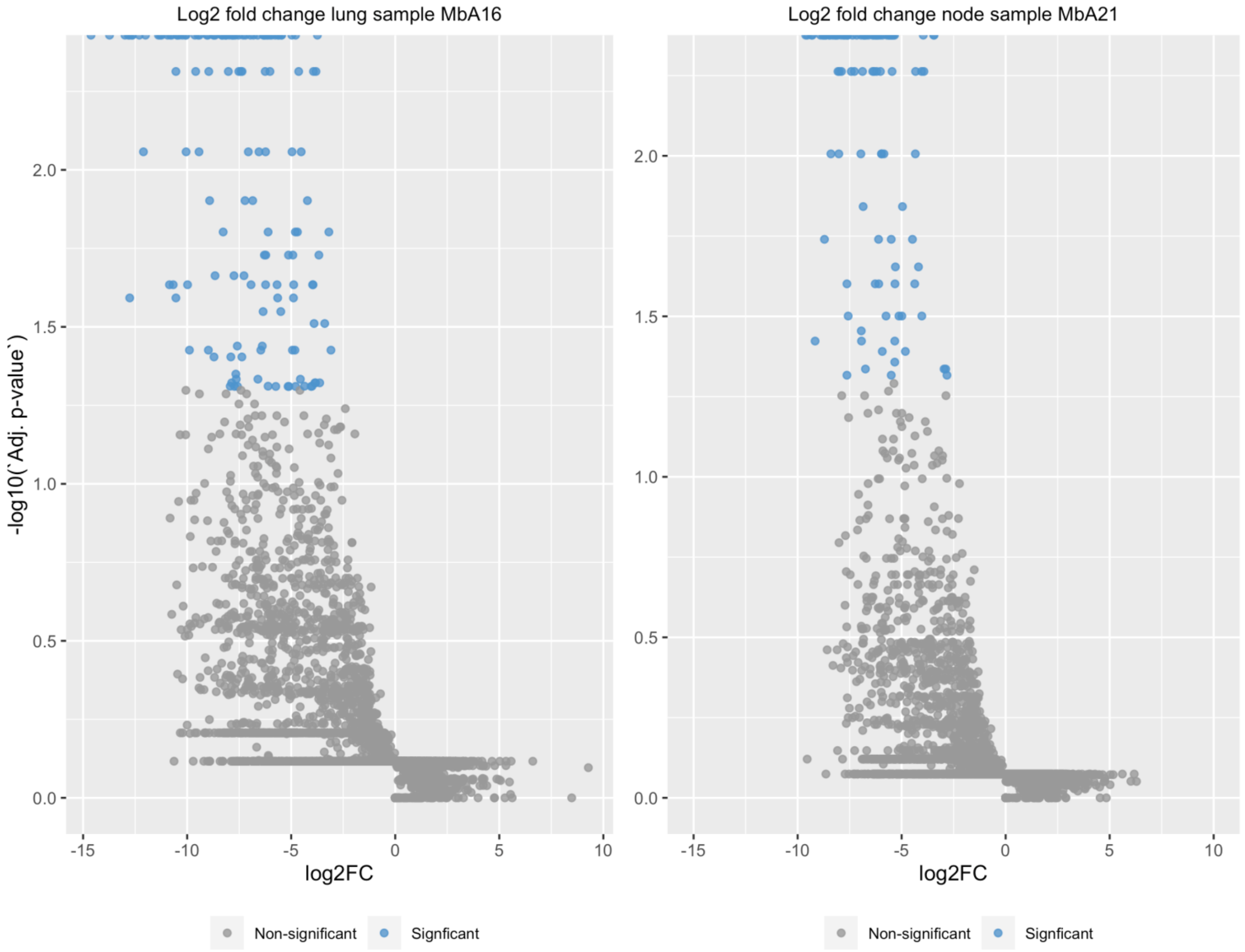


(A) Lung (B) Lymph node

**Figure S3. Volcano plots showing the distribution of $\log_2$ fold-changes and $-\log_{10}$ of adjusted p-values for representative lung (A) and lymph node (B) samples.** Adjusted p-values (BH-fdr correction) < 0.000001 cluster at the limits of the plot and precision reflects the number of resampling iterations (10,000)

**Figure S4**

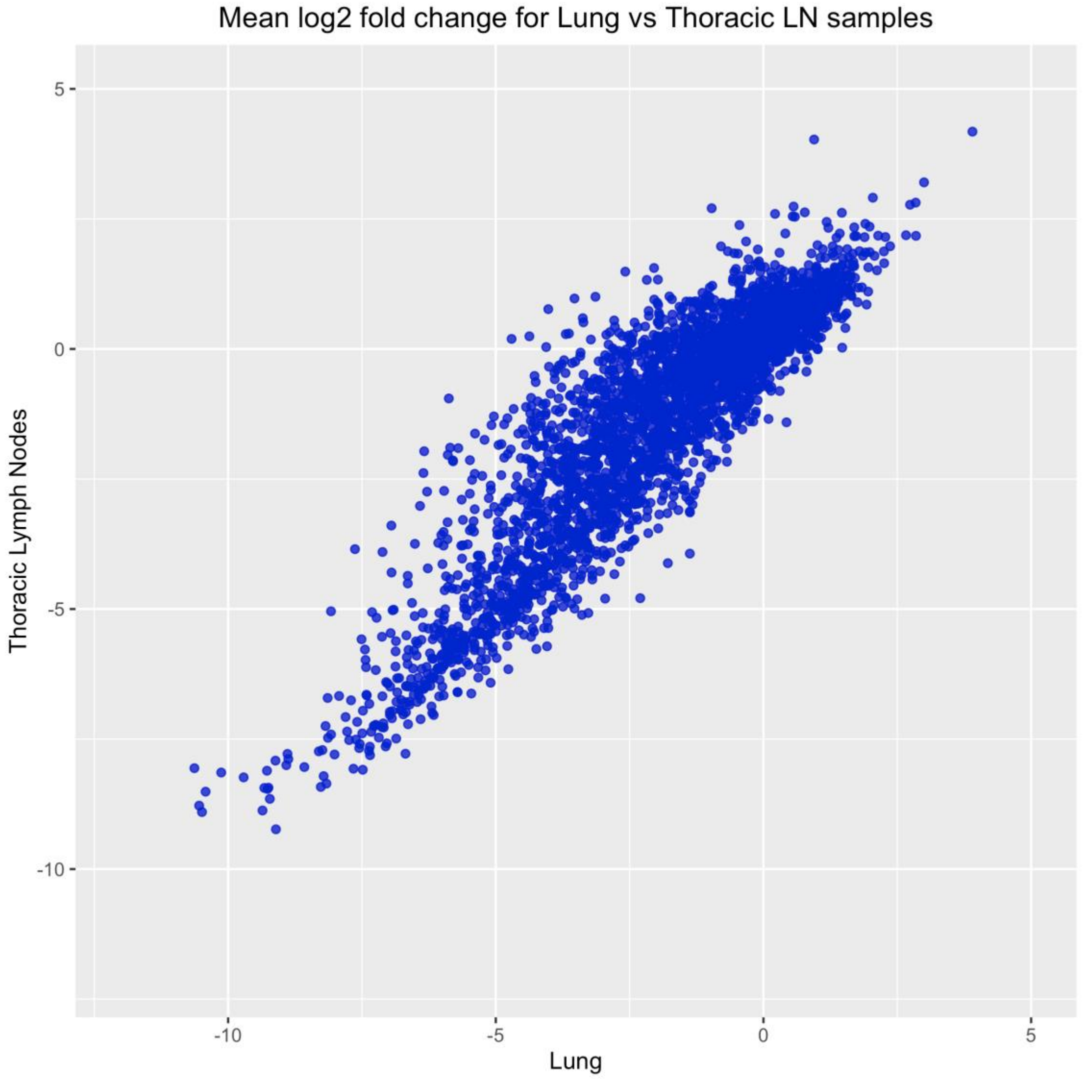


**Figure S4. Scatterplot of mean $\log_2$ fold change per gene for all lung samples against all thoracic lymph node samples**. Correlation between mean $\log_2$ fold change among genes between the tissues was calculated with Spearman's ranked correlation, $\rho$= 0.878, p-value < 2.2e-16.

**Figure S5**

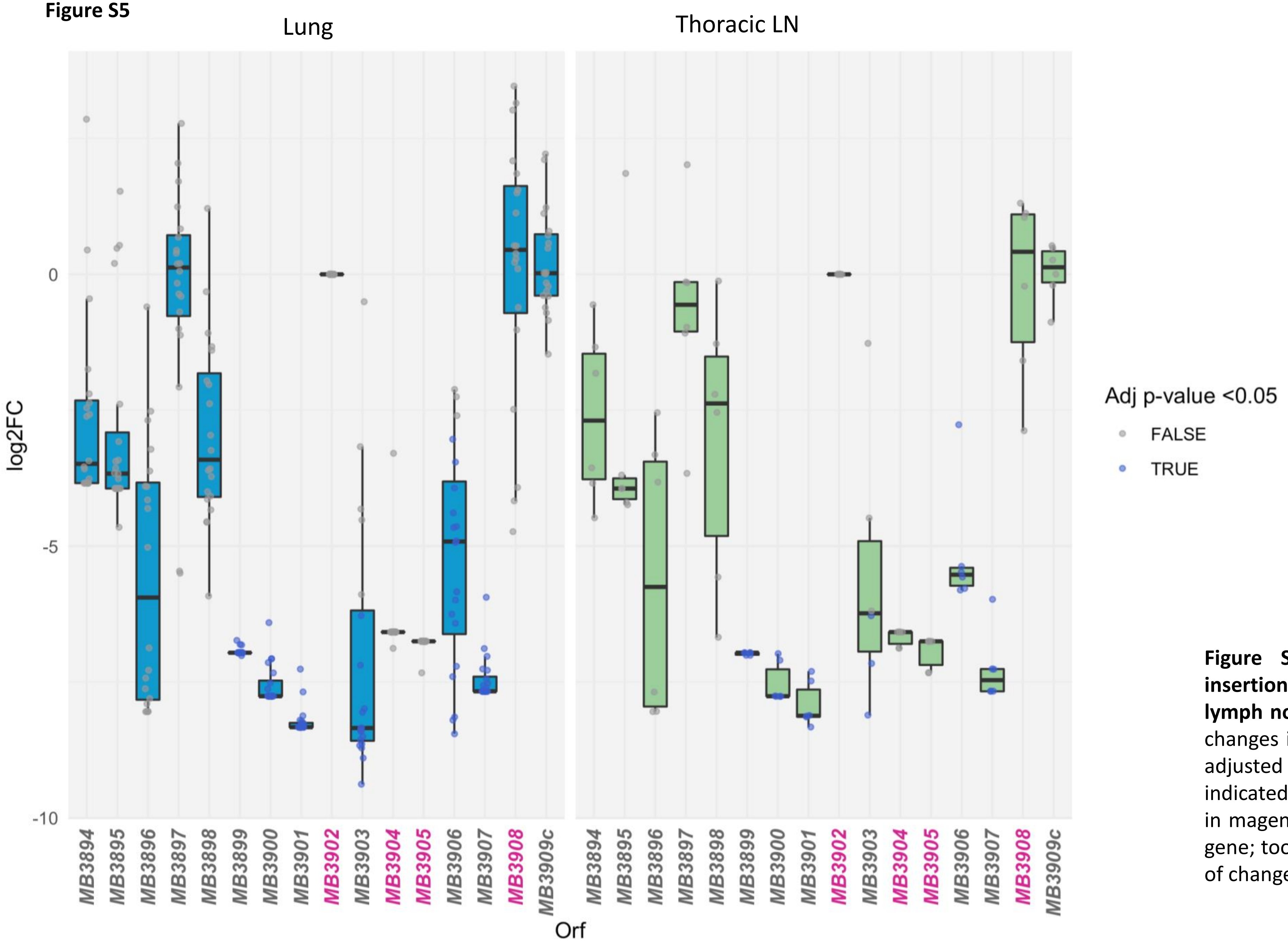


**Figure S5. Fold-changes caused by transposon insertions in *RD1*$^{BCG}$ and *RD1*$^{MIC}$ in the lungs and lymph nodes of infected cattle.** Boxplot for $log_2$ fold-changes in genes of the RD1$^{BCG}$ region. Samples with adjusted p-values (BH-fdr corrected) <0.05 are indicated with purple points. Gene names highlighted in magenta have fewer than 5 TA sites located in the gene; too few to determine the statistical significance of changes in insertion levels with this method.
